# The NIN-LIKE PROTEIN 7 (NLP7) transcription factor modulates auxin pathways to regulate root cap development

**DOI:** 10.1101/2022.03.12.484102

**Authors:** Narender Kumar, Chloe Caldwell, Anjali S. Iyer-Pascuzzi

**Author notes:** Author contributions: AIP conceived the project, experiments, and research plans; AIP and NK supervised the experiments; NK performed the experiments and analyzed the data; CC provided technical assistance to NK; AIP and NK wrote the article; AIP agrees to serve as the author responsible for contact and ensures communication. Funding information: This work was funded by the National Science Foundation grant number 1656392 to AIP.

## Abstract

The root cap surrounds the root tip and promotes root growth by protecting the root apical meristem, influencing root branching, and sensing environmental signals like nitrate. The root cap maintains a constant size through the coordination of cell production in the root meristem with cell release at the tip of the root, a process that requires an auxin minima in the last layer of the root cap. To perform its functions, the root cap must maintain a constant size and synchronize external cues with development, but mechanisms underlying such coordination are not well understood. Mutations in the NIN LIKE PROTEIN 7 (NLP7) transcription factor, a master regulator of nitrate signaling, lead to defects in root cap cell release and cell production. Nitrate impacts root development through crosstalk with auxin. Therefore, we hypothesized that NLP7 regulates root cap cell release and cell production by modulating auxin pathways. Here we show that mutations in NLP7 abolish the auxin minima required for root cap cell release and alter root cap expression levels of the auxin carriers PIN-LIKES 3 (PILS3) and PIN-FORMED 7 (PIN7). We find that NLP7 is required for proper root cap cell production and differentiation and for expression of transcription factors that regulate these processes. Nitrate deficiency impacts auxin pathways in the last layer of the root cap, and this is mediated in part by NLP7. Together, our data suggest that NLP7 integrates nitrate signaling with auxin pathways to optimize root cap development in response to external nitrate cues.

**One sentence summary:** The nitrate master regulator NLP7 controls root cap development through auxin pathways.

## Introduction

The root cap is a small but critical tissue at the tip of the root of angiosperms, gymnosperms and pteridophytes (Barlow, 2002) which functions to sense environmental cues, protect the Root Apical Meristem (RAM), influence root architecture (Xuan et al., 2015; Di Mambro et al., 2019), and defend against biotic and abiotic stress (Barlow, 2002; Kumpf and Nowack, 2015; Hawes et al., 2016; Driouich et al., 2021). In addition to its importance for plant health, the root cap is an excellent model to study developmental questions related to organ development. The tissue consists of two cell types: a lateral root cap that surrounds the root meristem and a columella root cap located at the root tip. In Arabidopsis, root cap cells in the lateral root cap undergo programmed cell death near the end of the meristematic region (Fendrych et al., 2014; Xuan et al., 2016; Huysmans et al., 2018). However, root cap cells in the last layer of the cap at the root tip are released as a single intact layer of living cells in a developmentally programmed manner (Vicré et al., 2005; Durand et al., 2009; Bennett et al., 2010). New cell production from columella stem cells (CSCs) compensates for release of the last layer. The coordination between root cap cell production and release is a key factor in maintaining root cap size (Dubreuil et al., 2018; Shi et al., 2018), which is essential for preserving root cap functions (Barlow, 2002; Roué et al., 2020). As it lies at the interface of the root and soil, the root cap must coordinate size homeostasis with environmental cues yet how it does so is not clear.

Auxin is a critical instructive signal for many plant developmental processes (Zhao, 2018; Gomes and Scortecci, 2021), including those in the root cap. The balance between the developmentally programmed release of outer root cap layers and the production of new columella cells is mediated in part through an auxin gradient in the columella and small peptide signaling pairs (Dubreuil et al., 2018; Shi et al., 2018). The root cap auxin gradient has a maxima in the quiescent center (QC) cells in the stem cell niche of the root apical meristem, and a minima in the separating layers at the root tip (Dubreuil et al., 2018). Recent work revealed that the auxin minima in the last layer is required for root cap cell release (Dubreuil et al., 2018). In addition to root cap cell release, auxin also impacts columella root cap cell production and differentiation. Auxin flow within the columella root cap is maintained in part through PIN-FORMED (PIN) auxin efflux carriers (PINs) (Blilou et al., 2005; Křeček et al., 2009). Mutations in PIN3/4/7 impact columella stem cell division and differentiation (Ding and Friml, 2010; Qin et al., 2019). Additionally, exogenous auxin treatment or auxin production within CSCs promotes their differentiation (Ding and Friml, 2010). This is mediated largely through the influence of auxin on the expression of *WOX5*, a transcription factor that is expressed in the QC and maintains CSCs in their undifferentiated state (Sarkar et al., 2007). The auxin response factors ARF10 and ARF16 also promote CSC differentiation (Want et al. 2005). Auxin pathways are thus essential to root cap development.

We recently found that NIN LIKE PROTEIN7 (NLP7), a plant-specific transcription factor, regulates the release of the last layer of root cap cells in Arabidopsis (Karve et al., 2016). In *nlp7-1*, cells in the last layer of the root cap are released as single cells instead of an intact layer, and the columella has fewer cell layers compared to wild type (Karve et al., 2016). Our work revealed that NLP7 modifies the expression of cell wall degrading enzymes and genes involved in root cap development, but how it impacted both root cap cell release and cell production was not clear.

NLP7 is well described as a major regulator of nitrate signaling (Castaings et al., 2009; Marchive et al., 2013; Guan et al., 2017; Zhao et al., 2018b). Nitrate assimilation and signaling have significant crosstalk with auxin transport and signaling pathways in the control of root development (Vega et al., 2019; Asim et al., 2020; Liu and von Wirén, 2022). For example, nitrate regulates root growth in part by modulating the phosphorylation status of the PIN2 auxin transporter (Otvos et al. 2021; Vega et al. 2021). Nitrate also induces the expression of the auxin receptor AUXIN SIGNALING F-BOX3 (AFB3) (Vidal et al., 2010; Vidal et al., 2013). The nitrate transceptor NRT1.1/NPF6.3 senses nitrate, transports both nitrate and auxin, and regulates auxin biosynthesis (Krouk et al., 2010; Mounier et al., 2014; Maghiaoui et al., 2020). Additionally, low nitrate conditions promote differentiation of CSCs through auxin pathways and *WOX5* (Wang et al., 2019). Several lines of evidence suggest that NLP7 may also play a role in auxin pathways. Studies of NLP7 genome wide binding sites and RNA-seq have identified auxin carriers PIN3 (O’Malley et al., 2016), PIN7 and PIN LIKES 3 (PILS3) (Marchive et al., 2013; Liu et al., 2017; Alvarez et al., 2020), the auxin receptor TIR1 (Marchive et al., 2013) and several auxin signaling proteins (Marchive et al., 2013; Alvarez et al., 2020) as putative direct targets of NLP7. ChIP and EMSA analyses revealed that NLP7 also binds to the promoter and activates expression of the auxin biosynthesis gene *TRYPTOPHAN AMINOTRANSFERASE RELATED 2 (TAR2)* (Zhang et al., 2021).

Given the role of both NLP7 and auxin in root cap development, and the relationship between nitrate and auxin pathways, we hypothesized that 1) NLP7 affects root cap size homeostasis by modulating auxin pathways in the root cap, and 2) NLP7 mediates the effect of nitrate on auxin pathways in the root cap. We found that auxin transporters have significantly decreased expression in the *nlp7-1* root cap and that auxin activates expression of *NLP7*. Our data show that the auxin minima in the last layer of the root cap, which is required for root cap cell release, does not occur in the *nlp7-1* mutant. Consistent with the root cap defects, we found that *nlp7-1* has decreased *WOX5* expression in the QC and increased differentiation of CSCs compared to wild type. We find that NLP7 mediates many of the effects of nitrate on auxin root cap pathways. Together, our data show that NLP7 links nitrate signaling with root cap cell production and release by modulating auxin pathways and expression of critical transcription factors, thereby enabling root growth under optimal conditions.

## Results

### Expression of auxin transporters is decreased in the root cap of *nlp7-1*

Columella-localized auxin efflux transporters PIN3, 4, and 7 mediate polar auxin transport in the root cap (Blilou et al., 2005; Křeček et al., 2009). Previous studies using DAP-seq (O’Malley et al., 2016) and ChIP-seq (Alvarez et al., 2020), showed that NLP7 binds to the upstream sequences of PIN3 (O’Malley et al., 2016) and PIN7 (Alvarez et al., 2020), and RNA-seq analyses revealed that PIN7 is highly upregulated in mesophyll protoplasts expressing NLP7 (Liu et al., 2017). Given these data, we hypothesized that NLP7 regulates auxin transport in the root cap by controlling the localization and expression of auxin carriers. We reasoned that deficiencies in auxin transport may play a role in the altered *nlp7-1* root cap phenotypes previously observed (Karve et al. 2016). We examined the spatial expression domain and level of PIN3 and PIN7 proteins in *nlp7-1* plants. Compared to wild type plants, the fluorescence intensity and spatial expression domain of *pPIN7:PIN7:GFP* was significantly decreased in the columella cells of *nlp7-1* plants (Figure 1A, B, E); however, the expression of PIN3 did not significantly change (Supplemental Figure S1).

**Figure 1.**
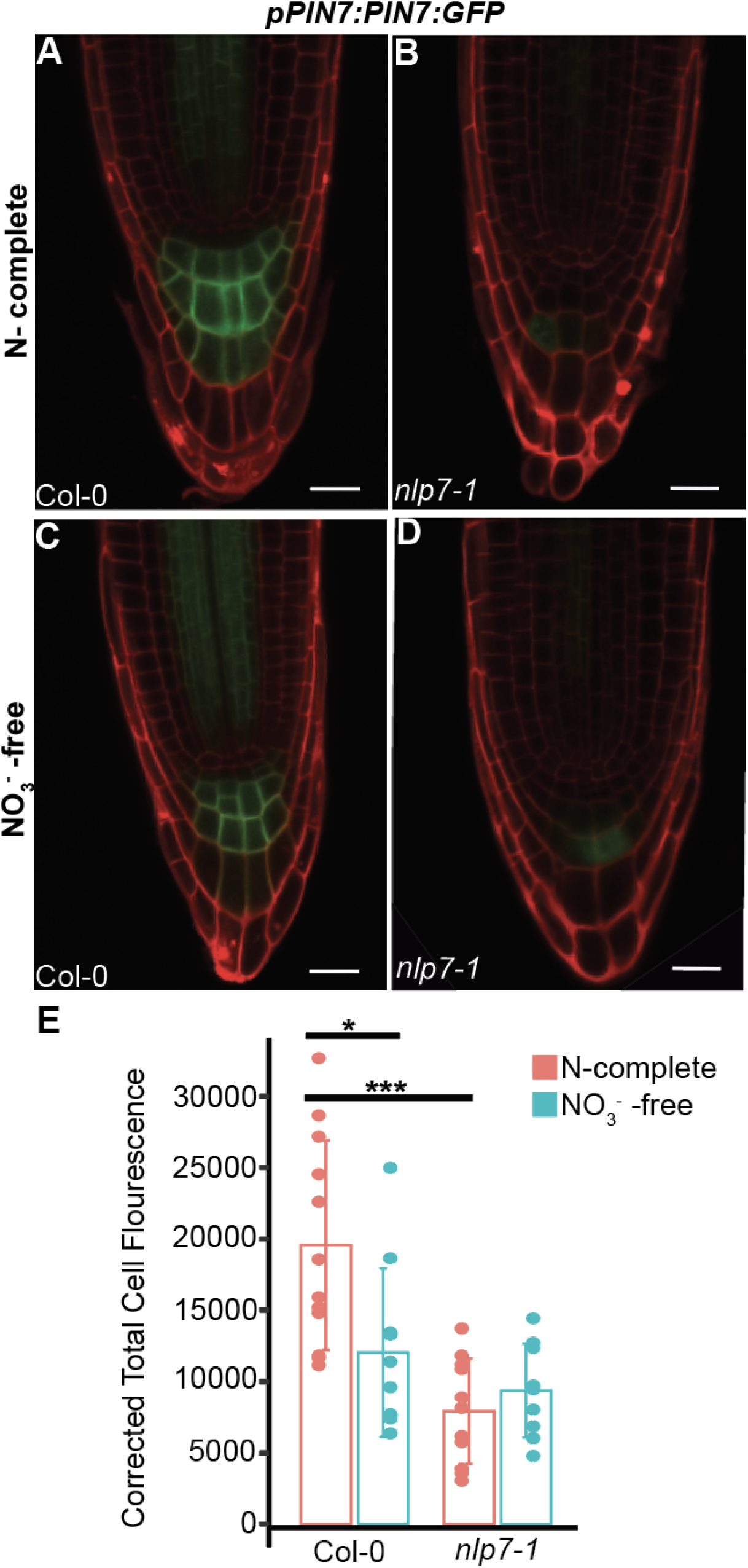
NLP7 impacts expression of auxin carriers in the root cap. **A - D.** Translational fusion (*pPIN7:PIN7:GFP*) of auxin carrier PIN7 in Col-0 and *nlp7-1* plant root cap of five-day-old plants grown on N-complete (A, B) and NO_3_^-^ - free media (C, D). **B.** Quantification of GFP fluorescence. PIN7 expression is significantly decreased in *nlp7-1*. Dots indicate the number of plants (n >10) used for each genotype and treatment. Three independent biological replicates were performed; a representative replicate is shown. Scale bar is 20 μm, magnification is 40X Asterisks indicate a significant difference with a two-tailed t-test (*P < 0.05, ***P < 0.001).

Nitrate regulates the subcellular localization pattern of the PIN2 auxin transporter in roots (Ötvös et al., 2021). To further investigate the relationship between auxin transporters, NLP7 and nitrate (NO_3_^-^), we grew wild type Col-0 and *nlp7-1* plants expressing *pPIN7:PIN7:GFP* on NO_3_^-^-free media. When grown on NO_3_^-^-free media, expression of PIN7 significantly decreased in wild type plants compared to growth on nitrogen (N)-complete media (Figure 1A, C, E). In contrast, expression of PIN7 in the root cap of the *nlp7-1* mutant did not significantly differ between *nlp7-1* plants grown on N-complete and NO_3_^-^ -free media (Figure 1B, D, E).

PIN LIKES (PILS) proteins are auxin carriers that reside in the endoplasmic reticulum to regulate intracellular auxin homeostasis (Barbez et al., 2012). Based on ChIP-chip and ChIP-seq data, *PILS3* is also a direct target of NLP7 (Marchive et al., 2013; Alvarez et al., 2020). *PILS3* is expressed predominantly in the lower layers of the root cap (Figure 2). Examination of *pPILS3:PILS3:GFP* expression in the *nlp7-1* mutant revealed that compared to wild type, PILS3 was significantly decreased in the root cap of *nlp7-1* (Figure 2). Growth of *pPILS3:PILS3:GFP/Col-0* plants on NO_3_^-^ -free media revealed that expression of PILS3 was also reduced in conditions lacking NO_3_^-^ (Supplemental Figure S2).

**Figure 2.**
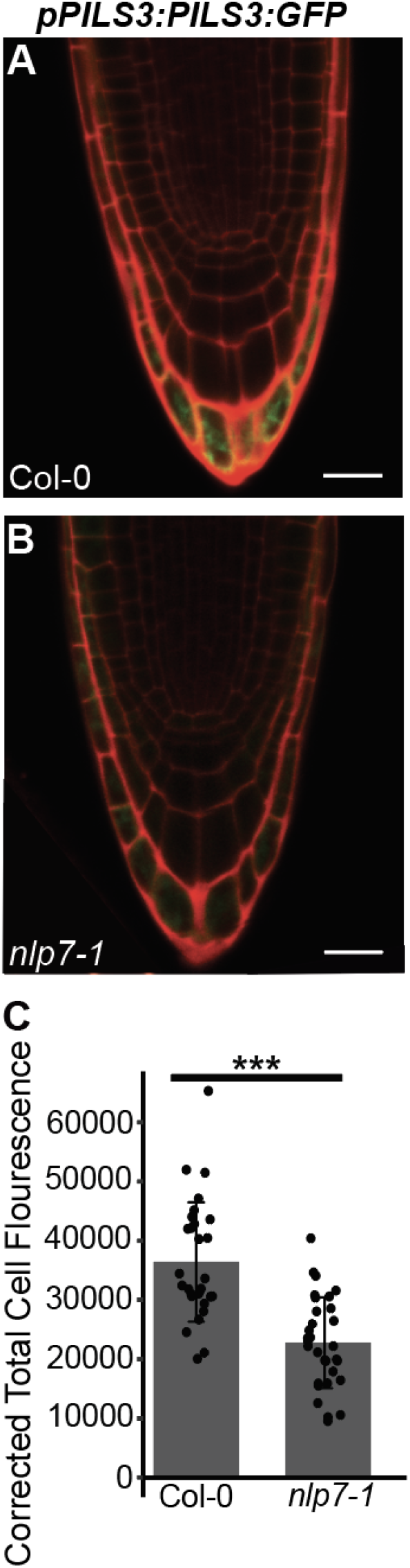
Expression of PILS3 is decreased in the *nlp7-1* mutant root cap compared to wild type Col-0. **A and B.** Translational fusion of *pPILS3:PILS3:GFP* in Col-0 (A) and *nlp7-1* (B) root cap of five-day-old plants. Scale bar is 20 μm, magnification is 40X. **C.** Quantification of GFP fluorescence. Black dots indicate individual plants. Two independent biological replicates were performed for a total of 30 roots per treatment; all are shown. Asterisks indicates a significant difference with a two-tailed t-test (*** = P < 0.001).

The reduced expression of PIN7 and PILS3 in the *nlp7-1* root cap suggests that NLP7 may impact auxin pathways in the root cap by modifying the expression of auxin carriers.

### Auxin responses increase in the last layer of the root cap of *nlp7-1*

The decrease in PILS3 expression in the *nlp7-1* root cap led us to question whether the auxin minima necessary for root cap cell release was altered in *nlp7-1*. To gain further insight into this, we generated an *nlp7-1* mutant expressing green fluorescent protein (GFP) from the synthetic auxin-responsive promoter (*DR5*). This auxin-responsive reporter revealed increased GFP intensity in the last layer of the *nlp7-1* root cap in 5 day old seedlings compared to wild type (Col-0) plants (Figure 3A, B, E). Notably, the increased *DR5:GFP* expression in *nlp7-1* was only observed in the last layer of the root cap, as auxin responses in upper tiers of the columella remained unchanged in *nlp7-1* compared to wild type (Supplemental Figure 3). Given the cross-talk between auxin and nitrate, we next examined whether the increased auxin response in the last layer of the *nlp7-1* root cap was impacted by nitrate. In wild type plants, auxin responses (as quantified by *DR5:GFP*) in the last layer of the 5 day old root cap were not significantly reduced when plants were grown on NO_3_^-^ -free media compared to N-complete media (Figure 3A, C, E). In *nlp7-1*, auxin responses in the last layer remained significantly higher than wild type in both conditions, and were not significantly different on either N-complete or NO_3_^-^ -free media (Figure 3B, D, E). Thus, regardless of nitrate, an auxin minima does not form in the last layer of the *nlp7-1* root cap. This lack of auxin minima could contribute to the aberrant root cap cell release in *nlp7-1*.

**Figure 3.**
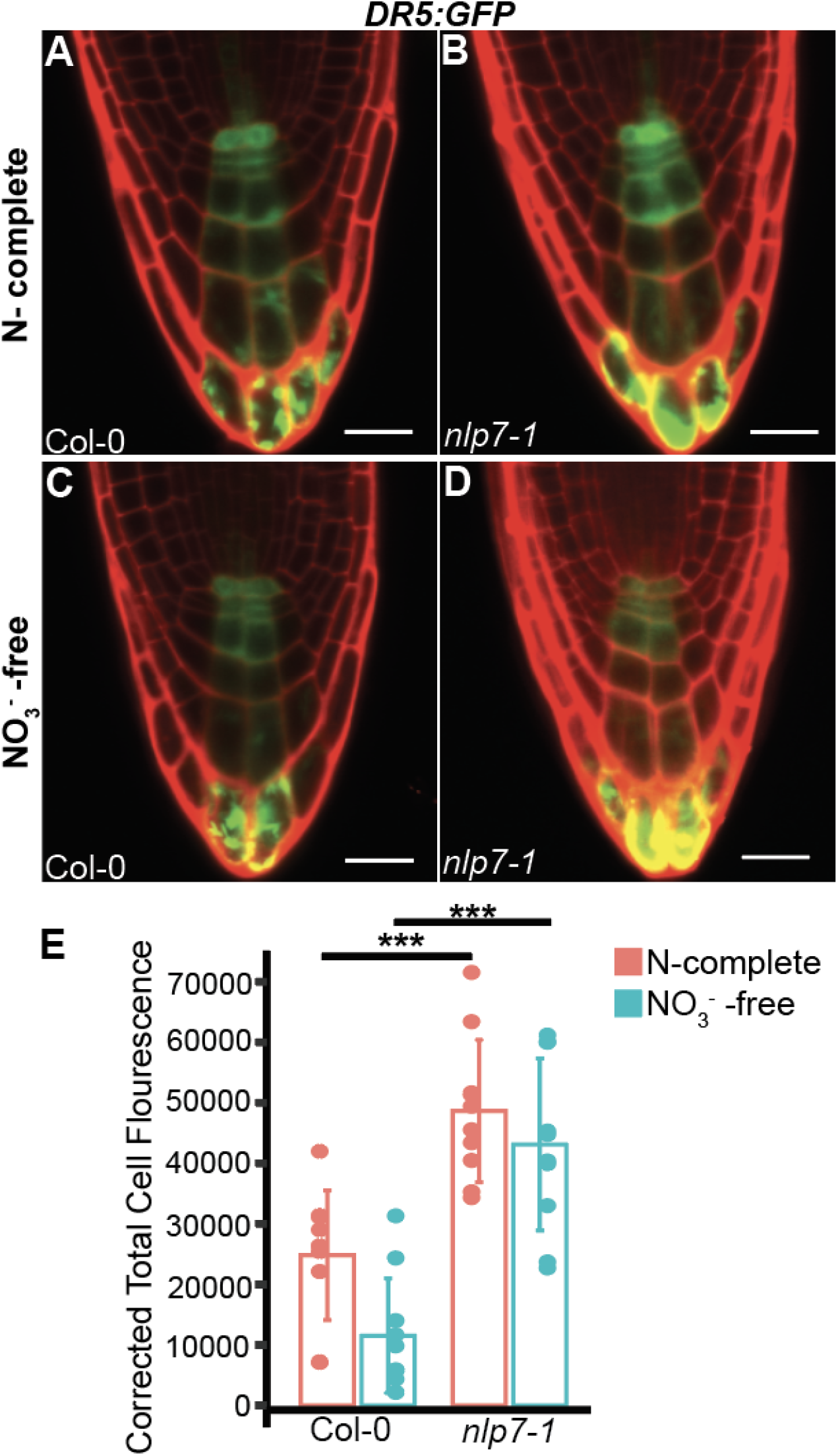
Auxin responses are increased in the last layer of the *nlp7-1* plant root cap. **A-D**. Auxin responsive synthetic promoter (*DR5:GFP*) expressed in the root cap of five-day-old Col-0 and *nlp7-1* plants grown in N-complete (A, B) and NO_3_^-^ - free conditions (C and D). **E.** GFP fluorescence intensity shows significant increase in the last layer of the *nlp7-1* root cap compared to wild type Col-0. Black dots indicate the number of plants (n = 10) used for each genotype and treatment. Three independent biological replicates were performed. A representative replicate is shown. Scale bar is 20 μm, magnification is 40X. Asterisks indicate a significant difference (***P < 0.001) using a two-tailed t-test.

### Auxin induces *NLP7* gene expression in the root cap

Our results reveal a role of NLP7 in modulating expression of PIN and PILS auxin carriers in the root cap. PIN auxin carriers are induced by auxin (Vieten et al., 2005; Benjamins and Scheres, 2008), and we questioned whether NLP7 may also be induced by auxin. Using the PLACE database of plant *cis-acting* regulatory elements (Higo et al., 1998; Higo et al., 1999), we identified three auxin-responsive cis-regulatory sequences (TGA-element, TGA-box, AuxRE) in the 3 kb region upstream of the NLP7 transcriptional start site. To test whether auxin (1-naphthaleneacetic acid, NAA) impacted *NLP7* expression, we grew an *NLP7* transcriptional reporter line (*pNLP7:GUS*) containing 3 kb upstream promoter sequence on MS media. Four day old seedlings were transferred either to control MS or MS + 0.3 μM auxin (NAA) supplemented plates for 24 hours. Root tips of *pNLP7:GUS* seedlings exposed to NAA showed enhanced *NLP7* expression (Figure 4).

**Figure 4:**
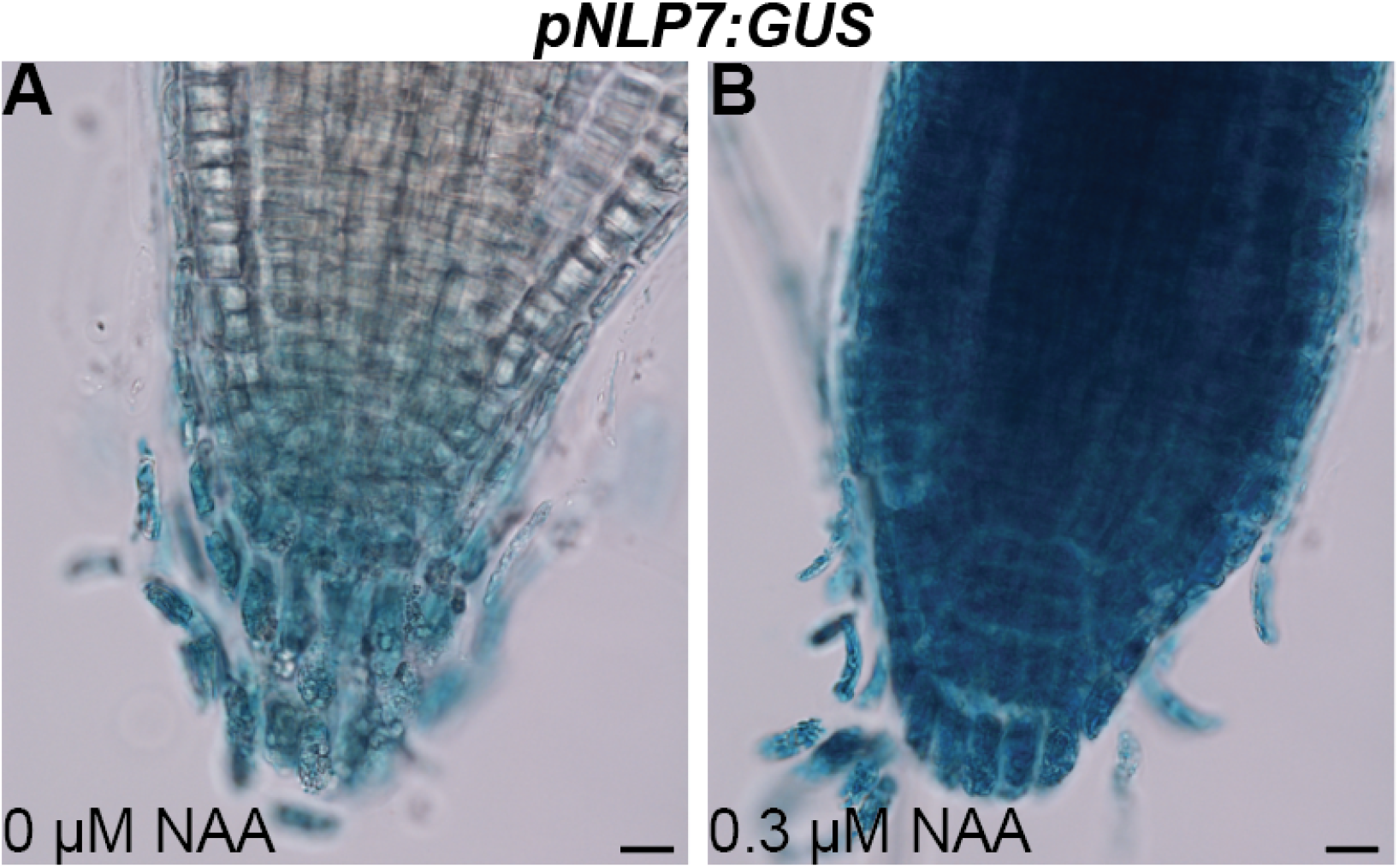
Visualization of *NLP7* expression in the root cap of wild type Col-0 using histochemical localization of GUS activity in *pNLP7:GUS* plants. A. Plant was grown on N-complete media without NAA. B. Plant was grown on N-complete media with 0.3 μM NAA. Scale bar is 20 μm, magnification is 60X.

### Increased CSC differentiation and less mitotically active columella cells are observed in the *nlp7-1* root cap

In addition to modified root cap cell release, the *nlp7-1* mutant also has fewer columella layers in the root cap compared to wild type plants (Karve et al., 2016). Given that the coordination between root cap cell release and stem cell division maintains the constant size of the root cap (Dubreuil et al., 2018; Shi et al., 2018), we hypothesized that NLP7 may impact columella cell production and differentiation. To examine the differentiation status of CSCs, we used Lugol’s stain, which detects starch granules that are present in differentiated columella cells (van den Berg et al., 1997). Approximately 20% of *nlp7-1* roots had no CSC layers, compared to 0% of wild type roots (Figure 5A-C).

**Figure 5.**
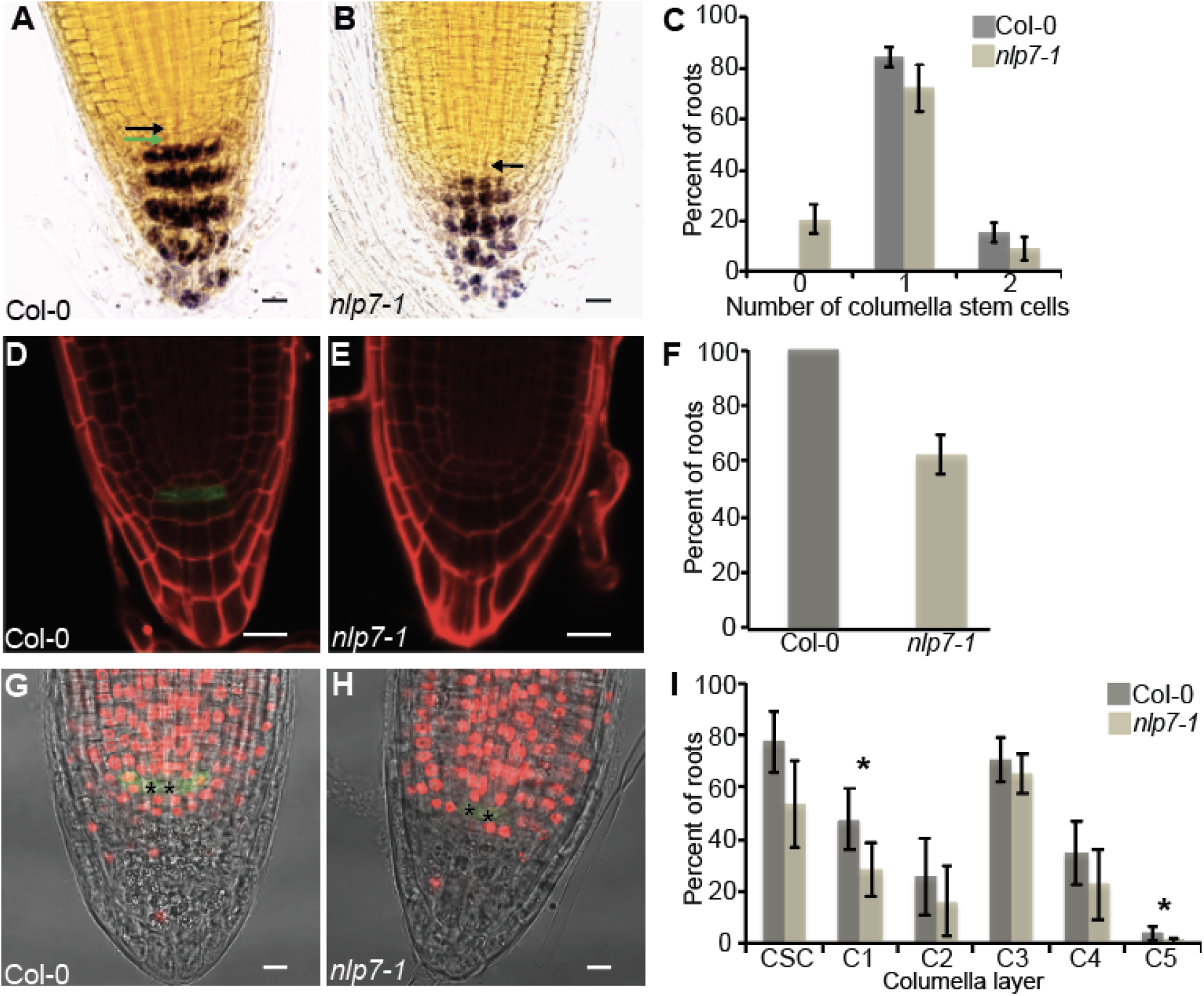
Columella cell differentiation is altered in *nlp7-1*.**A and B**. Lugol staining of the columella root cap of wild type Col-0 and *nlp7-1*. Black and green arrows indicate QC and columella stem cell, respectively. Columella stem cells are completely differentiated in some *nlp7-1* plants. Scale bar is 20 μm. **C.** Quantification of Lugol staining. Nearly 20% of *nlp7-1* roots have no columella stem cell layers (i.e. fully differentiated columella cells in the stem cell position). **D and E**. J2341 columella stem cell marker in Col-0 and *nlp7-1* plants. Scale bar is 20 μm; magnification is 40X. **F.** Percent of roots of wild type and *nlp7-1* that express J2341. Approximately 60%*nlp7-1* roots express the J2341 columella stem cell marker, in contrast to 100% of wild type plants. **G and H**. Columella cells are less mitotically active in the *nlp7-1* mutant. EdU staining of Col-0 and *nlp7-1* plants. Scale bar is 10 μm. **I.** Quantification of EdU staining. Columella cells are less mitotically active in *nlp7-1*. Asteriks indicate significant (P < 0.05) with a two-tailed t-test. For EdU and Lugol staining experiments more than 100 plants were examined. J2341 marker fluorescence was screened in more than 40 plants for each genotype. Representative images are shown; graphs reflect all plants examined.

To further investigate the effect of *nlp7-1* on stem cell activity, we examined expression of the J2341 enhancer trap line, which is expressed in CSCs in wild type plants. In *nlp7-1*, expression of J2341 was present in only ~ 60% of *nlp7-1* seedlings, compared to 100% of wild type seedlings (Figure 5D - F).

Columella stem cells divide continuously to produce cells that differentiate to mature columella cells. Lugol staining and J2341 marker data suggested that columella cell production may be changed in the *nlp7-1* mutant. To examine the mitotic status and division rate of CSCs, we used EdU staining. EdU is a thymidine analog that incorporates into newly synthesized DNA and produces fluorescence under the confocal microscope (Salic and Mitchison, 2008). A higher number of EdU stained cells indicates increased mitotic activity. This technique has been used to investigate the mitotic activity of CSCs (Hong et al., 2015). Four-day-old *nlp7-1* and wild type Col-0 seedlings were exposed to EdU-supplemented MS media plates for 24 hours. The *nlp7-1* mutant had a decreased percentage of EdU-stained columella cells in all columella layers, although the difference was only statistically significant in the C1 and C5 layers (Figure 5G-I). Together, these data reveal that the *nlp7-1* mutant has decreased columella stem cell layers, decreased mitotic activity in the columella, and an increase in CSC differentiation.

### WOX5 expression is decreased in the *nlp7-1* mutant

Our data suggest that NLP7 affects auxin transport in the root cap, as well as columella cell production and differentiation. The homeodomain transcription factor WOX5 maintains an auxin maxima at the QC by modulating auxin biosynthetic pathways (Tian et al., 2014; Savina et al., 2020), and mutations in *WOX5* lead to increased CSC differentiation (Sarkar et al., 2007). Expression of WOX5 is decreased in low nitrate conditions (Wang et al., 2019), suggesting a link between auxin and nitrate signaling in columella development. We hypothesized that the defects in columella cell production and differentiation in *nlp7-1* are partly due to changes in WOX5 expression. Examination of *pWOX5-GFP* in *nlp7-1* revealed that WOX5 expression was significantly reduced in *nlp7-1* mutant plants compared to wild type Col-0 plants (Figure 6A,B, E). The reduction in expression of WOX5 in *nlp7-1* is consistent with the increase in CSC differentiation observed in this mutant (Figure 5A-C). WOX5 expression is regulated by auxin response factors 10 and 16 (ARF10/16) (Wang et al., 2005), but we did not observe a difference in gene expression of ARF10/16 between whole roots of wild type and *nlp7-1* (Supplemental Figure S4).

**Figure 6.**
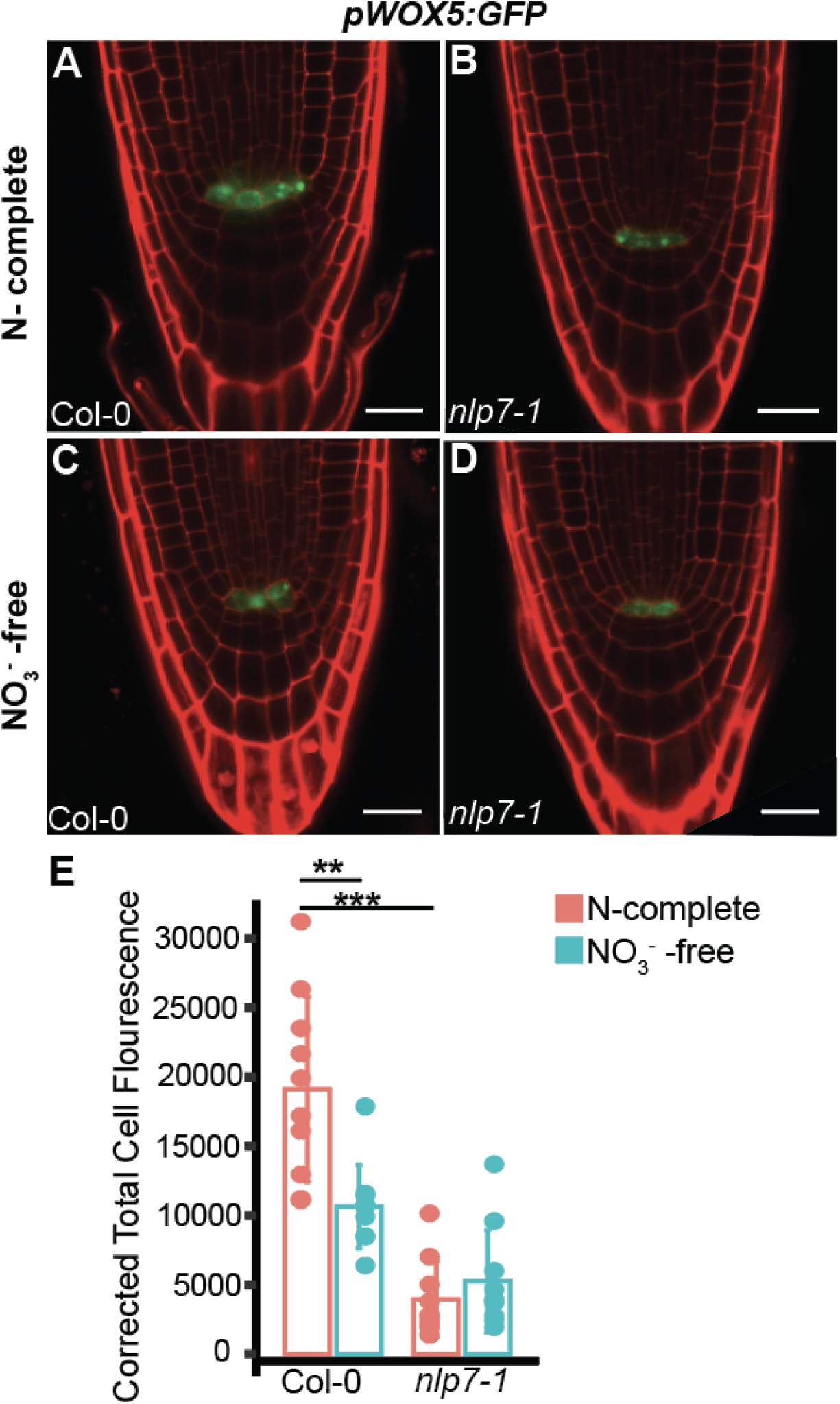
Expression of *pWOX5:GFP* is decreased in *nlp7-1*.**A - D.** *pWOX5:GFP* expressed in five-day-old Col-0 and *nlp7-1* plants grown on N-complete (A, B) and NO_3_^-^ - free media (C, D). **E.** Quantification of fluorescence intensity in A – D. Black dots indicate the plants (n = 10) used for each genotype and treatment. Three independent biological replicates were performed. A representative replicate is shown. Scale bar is 20 μm, magnification is 40X. Asterisks indicates a significant difference (**P < 0.01; ***P < 0.001) with a Wilcoxon Signed-Rank test.

To explore the link between NLP7, nitrate, and WOX5, we grew *pWOX5:GFP/* Col-0 and *pWOX5:GFP/nlp7-1* plants on NO_3_^-^ -free media. Consistent with previous work (Wang et al., 2019), in wild type plants, WOX5 expression decreased significantly on NO_3_^-^ -free media (Figure 6A, C, E). In contrast, WOX5 expression in *nlp7-1* plants was not significantly different on N-complete and NO_3_^-^ -free plates (Figure 6B, D, E). This suggests that the decreased WOX5 expression in wild type roots grown on NO_3_^-^ -free media is mediated by *NLP7*.

### PLETHORA 1 and 2 expression is decreased in *nlp7-1*

Our data show that NLP7 impacts the expression of auxin carriers PIN7 and PILS3 and the WOX5 transcription factor, suggesting that these defects contribute to the altered root cap phenotypes in *nlp7-1*. To further explore this, we examined an upstream regulator of WOX5. The APETALA2-type PLETHORA1 (PLT1) transcription factor, complexed with other proteins, directly binds to the promoter and induces the expression of *WOX5* (Shimotohno et al., 2018). PLT1/2 transcription factors and auxin work in a feedback loop to maintain the root stem cell niche (Aida et al., 2004; Blilou et al., 2005; Galinha et al., 2007; Mähönen et al., 2014; Santuari et al., 2016; Shimotohno et al., 2018). WOX5 and PLT1/2 also act downstream of auxin to maintain stem cell activity in the root meristem under low nitrate conditions (Wang et al., 2019). Examination of *pPLT1::PLT1:YFP* and *pPLT2::PLT2:YFP* in *nlp7-1* revealed that both PLT1 (Figure 7A -C) and PLT2 (Figure 8A-C) expression significantly decreased in *nlp7-1* compared to wild type plants. Notably, PLT2 directly targets and represses BEARSKIN2 (BRN2; Santuari et al., 2016), a NAC transcription factor that promotes root cap cell release (Bennett et al., 2010; Kamiya et al., 2016). We previously found that BRN2 is activated in *nlp7-1* compared to wild type (Karve et al., 2016). The repression of PLT2 in *nlp7-1* may contribute to this activation, and thus to the root cap cell release phenotype in *nlp7-1*.

**Figure 7.**
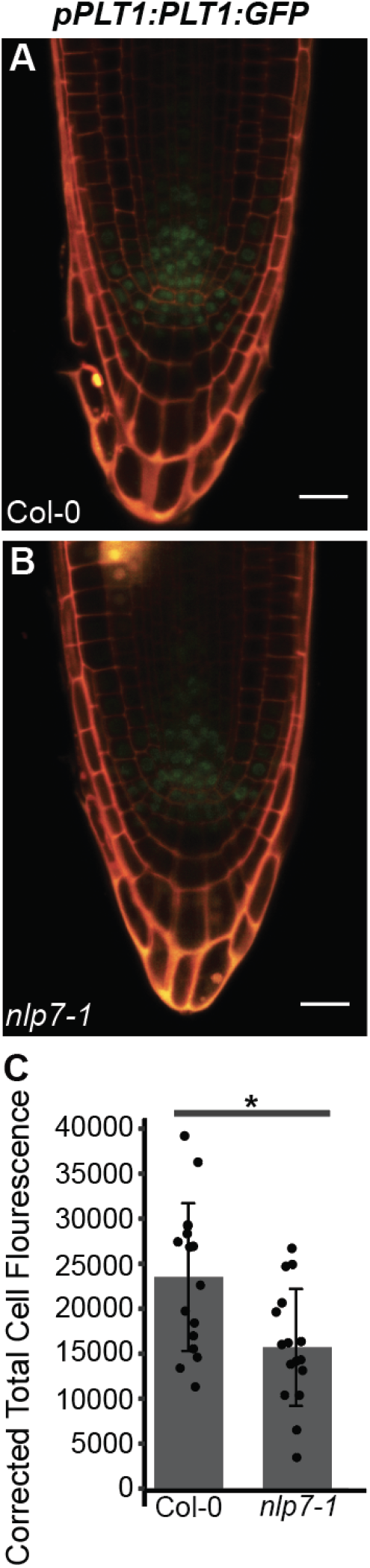
Expression of PLETHORA1 is reduced in *nlp7-1* compared to wild type plants. **A - B.** *pPLT1:PLT1:YFP* expressed in the root cap of five-day-old Col-0 (A) and *nlp7-1* (B) plants. **C.** Quantified fluorescence intensity (from A and B) shows a significant decrease in *nlp7-1* plants. Black dots indicate the plants (n = 16) used for each genotype and treatment. Three independent biological replicates were done. A representative replicate is shown. Scale bar is 20 μm, magnification is 40X. Asterisks indicates significant difference with a two-tailed t-test (*P < 0.05).

**Figure 8.**
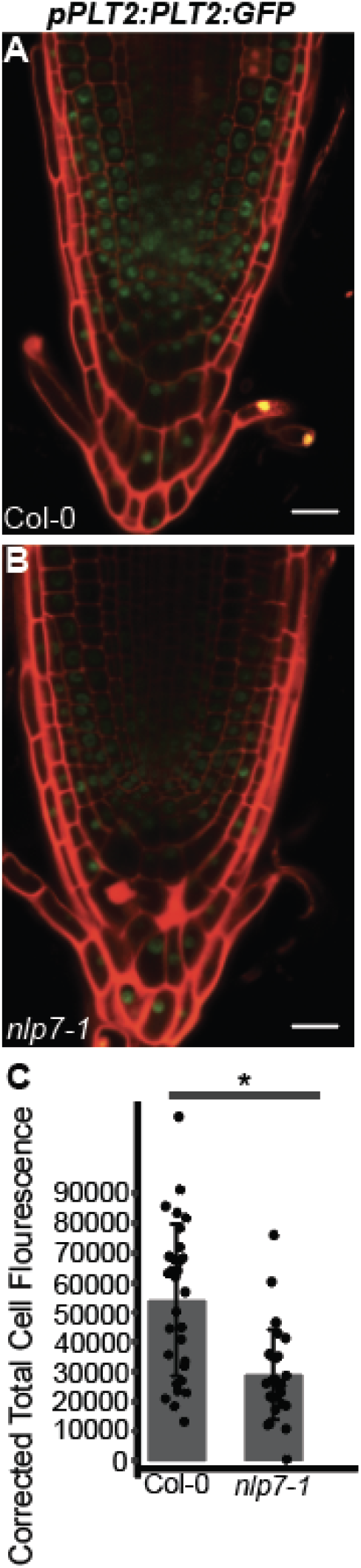
**A and B.** Expression of PLETHORA2 is reduced in *nlp7-1* compared to wild type plants. **A - B.** *pPLT2:PLT2:YFP* expression in Col-0 (A) and *nlp7-1* (B). **C.** Quantified fluorescence in A and B. Black dots indicate the plants (n = 30) used for each genotype and treatment. Three independent biological replicates were done; all replicates are shown. Scale bar is 20 μm, magnification is 40X. Asterisks indicates significant difference with the Wilcoxon Signed-Rank test (*P < 0.05).

### Model for NLP7 function in root cap development

Together, our data lead to a model in which NLP7 regulates root cap cell release and cell production by impacting the expression of auxin carriers and transcription factors critical for root cap development (Figure 9). *NLP7* is not transcriptionally regulated by NO_3_^-^ (Castaings et al., 2009), but in conditions with sufficient nitrate, NLP7 protein is retained in the nucleus (Marchive et al., 2013; Karve et al., 2016; Liu et al., 2017; Chu et al., 2021). In these conditions, in wild type plants, NLP7 promotes expression of PILS3, which reduces the nuclear auxin responses in the last layer of the root cap and may contribute to an auxin minima in the last layer of the root cap. This in turn promotes root cap cell release in NO_3_^-^-sufficient conditions. NLP7 also modulates expression of PLT1/2 and WOX5, thereby impacting columella cell production. Auxin in the root cap activates expression of *NLP7*, which likely reinforces the impact of NLP7 on root cap development. Thus, in conditions of sufficient NO_3_^-^, NLP7 promotes columella size homeostasis and continued function of the root cap.

**Figure 9.**
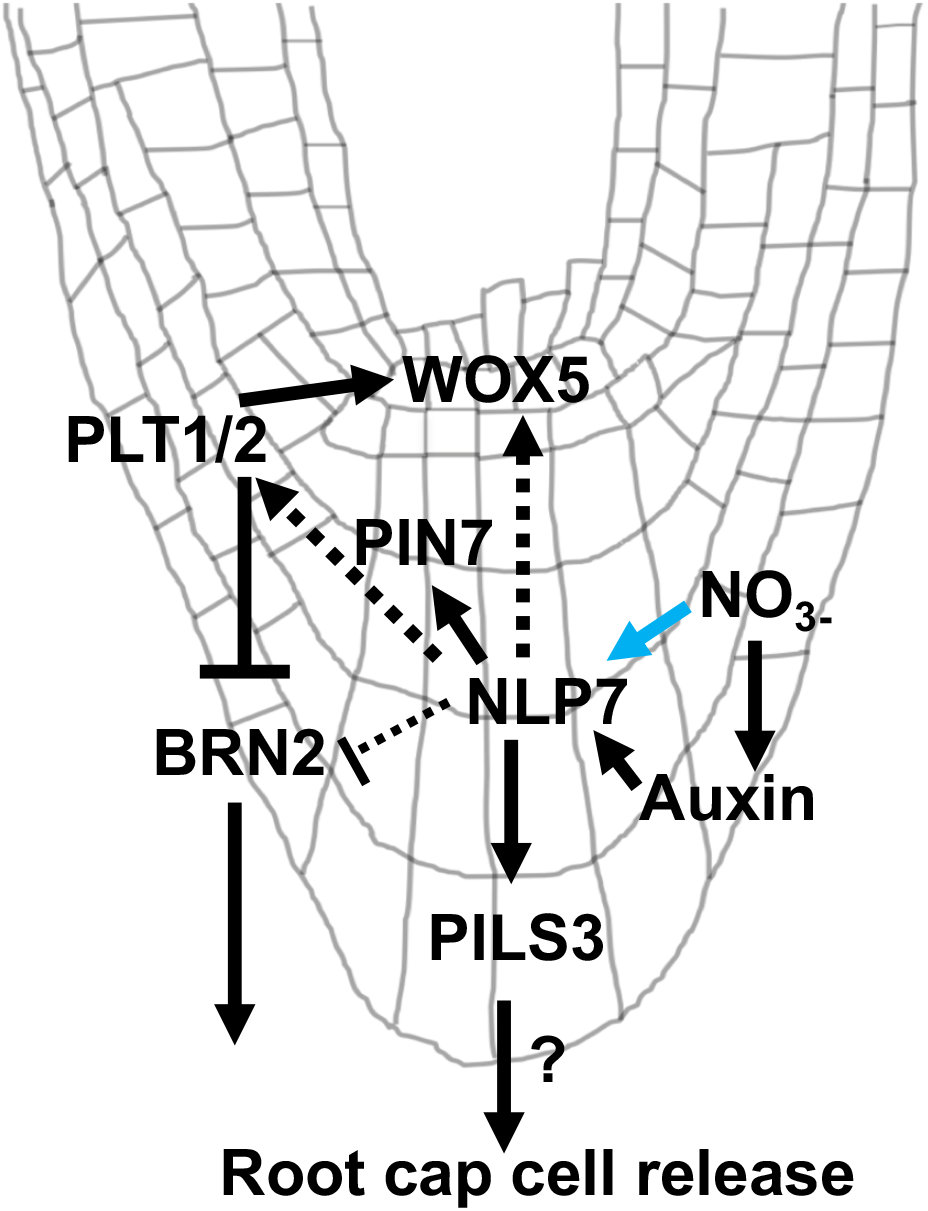
Model for NLP7 control of root cap cell release and production. Includes information obtained from studies of transcriptional regulation (i.e. transcription factor binding studies or change in gene expression level in a mutant) and post-translational regulation (protein expression in *nlp7-1*). Note that nitrate does not transcriptionally induce NLP7 but promotes its localization in the nucleus. Question mark indicates that the role of PILS3 in regulating root cap cell release is not clear. Solid and dashed lines indicate putative direct and indirect interactions, respectively.

## DISCUSSION

The root cap influences root meristem activity through the coordination of root cap cell release with cell production (Dubreuil et al., 2018; Shi et al., 2018), affects lateral root development and root architecture (Xuan et al., 2015; Di Mambro et al., 2019), protects the root tip from mechanical damage (Iijima et al., 2004; Fang et al., 2021) and soil microbes (Wen et al., 2009; Cannesan et al., 2012; Hawes et al., 2016; Karve et al., 2016; Tran et al., 2016; Driouich et al., 2021) and perceives environmental signals including nutrients (Arnaud et al., 2010; Kanno et al., 2016; He et al., 2021) and gravity (Strohm et al., 2012; Su et al., 2017). To perform these critical functions, the root cap must synchronize environmental signals with growth and developmental processes, but the mechanisms underlying such coordination are not well understood.

One of these environmental signals is nitrate. Nitrogen is a critical macronutrient for plant growth, and most crops absorb nitrate as their main nitrogen supply (Kant, 2018; Zhao et al., 2018a). Nitrate has major impacts on root development. Localized nitrate patches result in an increase in lateral root elongation and density in Arabidopsis (Linkohr et al., 2002; Sun et al., 2017; Liu et al., 2020). Moderate, homogenous nitrate levels promote primary and lateral root growth while high homogenous nitrate levels suppress growth (Asim et al., 2020; Boer et al., 2020; Liu and von Wirén, 2022). Nitrate mediates these functions largely through its impacts on endogenous auxin pathways. Numerous aspects of nitrate transport and signaling interact with auxin biosynthesis, signaling and transport (Vega et al., 2019; 2021; Asim et al., 2020; Otvos et al. 2021; Liu and von Wirén, 2022). Multiple studies have identified auxin-related genes as differentially expressed in response to nitrate (Gifford et al., 2008; Vidal et al., 2010; 2013). In the root cap, nitrate is required for auxin pathways (our study and (Wang et al., 2019)). The interaction between nitrate response and auxin pathways allows plants to coordinate nitrate availability with growth and development.

Much of the research which examines the relationship between nitrate and auxin has focused on the impact of nitrate on lateral root development and root architecture (reviewed in (Sun et al., 2017; Liu et al., 2020)). Here we show that NLP7, a master transcriptional regulator of nitrate signaling, regulates root cap cell release and production through auxin pathways. By integrating nitrate signals with plant developmental pathways in the root cap, NLP7 promotes root growth when nitrate supply is sufficient.

### NLP7 modulates expression of auxin carriers to alter root cap release

Mutations in NLP7 lead to the release of the last layer of cells as single cells, and fewer columella layers. We found that *nlp7-1* does not have an auxin minima in the last layer of the root cap, which likely contributes to the aberrant cellular release. The lack of auxin minima in *nlp7-1* appears to come in part from the impact of NLP7 on PILS3. Putative auxin carriers, PILS, are localized to the endoplasmic reticulum (ER), and control intracellular auxin homeostasis (Barbez et al., 2012). PILS efflux auxin out of the cytoplasm into the ER lumen. Less auxin is available to enter the nucleus, leading to decreased auxin responses. PILS3 as a putative direct target of NLP7 (Marchive et al., 2013; Alvarez et al., 2020). PILS3 is expressed in the lower tiers of the root cap, and PILS3 expression is decreased in *nlp7-1* compared to wild type. Decreased PILS3 expression in *nlp7-1* would result in less auxin influx into the ER and an increase in the available pool of nuclear auxin in the cell. This could promote the increased auxin responses observed in the last layer of the *nlp7* root cap.

NLP7 protein is present in the cytoplasm in nitrate-free conditions and is retained in the nucleus upon exposure to nitrate (Marchive et al., 2013; Liu et al., 2017; Chu et al., 2021). Consistent with cytoplasmic localization of NLP7 in nitrate-free conditions, in wild type plants under nitrate-free conditions, PILS3 expression is repressed (Supplemental Figure S2). However, an auxin minima in the last layer of the root cap still occurs in wild type plants in nitrate-free conditions. This suggests that NLP7 may also function in the cytoplasm to regulate auxin homeostasis in the last layer of the root cap in nitrate-free conditions.

Mutations in NLP7 also resulted in decreased expression of PIN7. PIN3, 4 and 7 are important for redirecting auxin to the lateral root cap and subsequently shootward. Due to their nonpolar localization on the plasma membrane, a directional auxin flux toward the root tip does not result from these transporters, and they are not thought to play a major role in the auxin concentration gradient in the root cap (Dubreuil et al., 2018). The impact of decreased PIN7 protein on the root cap phenotype in *nlp7-1* is not completely clear. Mutations in *PIN7* increase columella stem cell number (Ding and Friml, 2010). However, we observed an increase in columella cell differentiation in *nlp7-1*. Our results could be due to the decreased expression of WOX5 in *nlp7-1*, because *wox5* is epistatic to *pin7* (Ding and Friml, 2010).

The decreased expression of PIN7 in *nlp7-1* may be related to cell wall changes in *nlp7-1*. We previously showed that cell wall-related processes are impacted in *nlp7-1* (Karve et al., 2016). The *nlp7-1* mutant root cap has decreased homogalacturonan (HG), a major component of pectin, and elevated gene expression levels of several cell wall degrading enzymes (Karve et al., 2016). Gene expression of *CELLULASE5* (*CEL5*), which encodes a cellulase degrading enzyme required for root cap cell release (del Campillo et al., 2004), is upregulated in *nlp7-1*, and the *nlp7-1 cel5* double mutant has restored wild type root cap cell release (Karve et al., 2016). Since proper auxin transport is required for the cell wall remodeling which occurs during root cap cell separation (Dubreuil et al., 2018), the decreased expression of PIN7 in *nlp7-1* may contribute to the previously observed defects in cell walls. Alternatively, due to the redundancy in expression between PIN7 and PIN4, it is possible that decreased expression of PIN7 does not result in an observable phenotype in *nlp7-1*.

### NLP7 affects WOX5 expression and columella stem cell activity

The QC-localized transcription factor WOX5 is required to maintain columella stem cells in their undifferentiated state. We observed decreased expression of WOX5 and increase in CSC differentiation in *nlp7-1*. WOX5 protein expression decreased in wild type plants when grown on nitrate-free media compared to N-complete media, but remained unchanged in the *nlp7-1* mutant. This suggests that NLP7 mediates the effect of low nitrate on WOX5 and CSC differentiation. However, given that NLP7 and WOX5 are not expressed in the same spatial domain, the effect of NLP7 on WOX5 is likely indirect.

### NLP7 impacts additional transcription factors that determine root stem cell specification and cell release

PLT1/2 transcription factors maintain stem cell activity and the QC (Aida et al., 2004; Blilou et al., 2005; Galinha et al., 2007; Mähönen et al., 2014; Santuari et al., 2016; Shimotohno et al., 2018) and induce *WOX5* expression (Shimotohno et al 2018). PLTs are transcriptionally activated by auxin in the stem cell niche (Aida et al., 2004; Mähönen et al., 2014), and PLT2 regulates auxin biosynthesis and transport in the QC (Santuari et al., 2016). Previous work showed that PLT1/2 expression level in the QC decreases in low nitrate conditions (Wang et al., 2019). Consistent with this, we observed that the PLT1/2 protein level is reduced in *nlp7-1* compared to wild type plants.

PLT1 and NLP7 expression domains do not overlap, suggesting that an indirect interaction exists between them. Although the PLT2 and NLP7 expression domains overlap, neither is a transcriptional target of the other, based on genome-wide binding assays (Marchive et al., 2013; Santuari et al., 2016; Alvarez et al., 2020). Curiously, PLTs may function during the nitrate response. Among the genes repressed by PLT1, PLT2, and PLT7, the GO terms ‘nitrate transport’, ‘response to nitrate’, ‘nitrogen compound transport’ and ‘response to nitrogen’ compound are enriched (Santuari et al., 2016). NRT1.1 also is a putative direct target of PLT2 (Santuari et al., 2016). Both NLP7 and PLTs interact with the transcription factor teosinte branched 1/cycloidea/proliferating cell factor1-20 (TCP20; (Guan et al., 2017; Shimotohno et al., 2018)). PLT1/3 physically interact with TCP20 and other transcription factors to activate WOX5 gene expression (Shimotohno et al., 2018), while NLP7 forms heterodimers with TCP20 (Guan et al., 2017). Whether PLT2 and NLP7 interact, and whether this occurs through a TCP transcription factor, remains to be explored. Additionally, whether these proteins together impact WOX5 or nitrate-induced gene expression is unknown.

In conclusion, our data show that NLP7 impacts root cap cell release and cell production by modulating the expression of auxin carriers and root cap developmental regulators. Our data suggest that NLP7 modulates root cap development in response to external nitrate cues, thereby allowing plants to optimize root growth in different environments. Additional work to investigate the genetic and physical interactions among NLP7, WOX5, PINs, PILS, and PLT1/2 is needed to fully understand the molecular mechanisms underpinning the relationship between nitrate and root cap development.

## MATERIALS AND METHODS

### Plant material and growth conditions

All plants, except the enhancer trap line J2341 are in the Col-0 genetic background. J2341 (in the ecotype C24) was crossed to both Col-0 and *nlp7-1* to maintain comparable backgrounds. The *nlp7-1* homozygous mutant line (*At4g24020;* Salk_026134, as described in Castaings et al. 2009 and (Karve et al., 2016) was crossed with pDR5:GFP:GUS, pPIN7:PIN7:GFP, pPIN3:PIN3:GFP, pPILS3:PILS3:GFP, pWOX5:GFP, pPLT1:PLT1:YFP, and pPLT2:PLT2:YFP (all in Col-0) to produce translational and transcriptional fusion in *nlp7-1* homozygous background. All crosses were tested for *nlp7-1* homozygosity via PCR with *NLP7* gene-specific and T-DNA border primers. If GFP fluorescence was not observed, plants were tested via PCR with GFP-specific primers to examine whether the lack of fluorescence was due to lack of construct or the absence of NLP7. Seeds of *pPIN7:PIN7:GFP* were gifted from Jiri Friml,*pPIN3:PIN3:GFP* from Christopher Staiger, *pPLT1:PLT1:YFP* and *pPLT2:PLT2:YFP* from Ben Scheres, *pPILS3:PILS3:GFP* from Jürgen Kleine-Vehn, *pWOX5:GFP* from Philip Benfey, and *pNLP7:GUS* from Anne Krapp.

Seeds were surface sterilized with 50% bleach for five minutes, stratified at 4°C for two days, and subsequently planted on Murashige and Skoog (MS) basal media plates (MSP01, Caisson labs, Utah, USA) with 0.8% agar and allowed to grow vertically. For growth on nitrate-free media, MS without nitrogen (MSP21, Caisson labs, Utah, USA) was supplemented with 2.5 mM ammonium succinate as in (Konishi and Yanagisawa, 2013; Xu et al., 2016; Liu et al., 2017). All plants were grown at 23°C under a 16-h light/8-h dark cycle photoperiod.

### Lugol and EdU staining

Freshly prepared Lugol’s solution (Iodine crystal and Potassium Iodide) was diluted 1:5 with sterilized water. Five day old seedling root tips were stained for 10 seconds and mounted immediately onto a clean microscopic glass slide with chloral hydrate (chloral hydrate: glycerol: water in 8:3:1 ratio). Lugol stained root cap were imaged with an Olympus (DP80) under a 63X magnification microscope equipped with Olympus DP80 camera. Stained starched granules marked the differentiated columella layers in the root cap.

For EdU (5-ethyny-2’-deoxyuridine) staining, seeds were planted and germinated on MS media plates with 0.8% agar, and pH was adjusted to 5.7. Four day old seedlings were transferred to MS plates supplemented with 10μM EdU (Invitrogen Click-iT EdU imaging kit) and allowed to grow for 24 hours. Roots were cut and vacuum infiltrated for one hour in a freshly prepared fixative solution containing (3.7% (v/v) paraformaldehyde (Sigma) and 1% (v/v) Triton-X 100 (Sigma) in 1X PBS solution (Vivantis). After fixation, samples were washed twice with 3% (w/v) bovine serum albumin or BSA (Santa Cruz) in 1X PBS solution. Washed samples were incubated in 50 μl Click-iT^®^ reaction cocktail with 1 × Click-iT^®^ EdU reaction buffer (43 μl), CuSO_4_ (2μl), Alexa Fluor^®^ azide (0.12 μl), and 1 × Click-iT^®^ EdU buffer additive (5 μl), as in (Hong et al., 2015). Samples were incubated for one hour in dark at room temperature and subsequently washed again with 3% (w/v) bovine serum albumin or BSA (Santa Cruz) in 1X PBS solution. Samples were stored in 1X PBS solution until ready for imaging as described by (Hong et al., 2015). EdU stained root caps were mounted in 1X phosphate-buffered saline (PBS) solution and imaged using a LSM800 laser scanning confocal microscope (Zeiss, Jena, Germany) using 60X magnification. The number of columella layers was counted based on the EdU stained cells in the root cap. Images of EdU stained cells were captured using argon laser excitation at 647 nm and emission between 655 nm and 750 nm.

### Confocal Microscopy and Data Analysis

Five day old seedling roots were stained with 10 μg/mL propidium iodide and imaged with a LSM800 laser scanning confocal microscope (Zeiss, Jena, Germany) with the Argon laser set to an excitation of 488 nm and 505 to 550 nm emission filter for observation of GFP fluorescence. The intensity of the fluorescence signal was measured using ImageJ software (ImageJ 1.52q, (Schneider et al., 2012)) and corrected total cell fluorescence (CTCF) was calculated using the formula (CTCF = Integrated Density - (Area of selected cell X Mean fluorescence of background readings). All images for all samples within an experiment were captured using equal laser power, detector gain, and pinhole in the xy plane. Figures were generated in Adobe Illustrator. Confocal images were adjusted for brightness and contrast using Adobe Photoshop. Both wild type and mutant images were adjusted in the same way, and the entire image was adjusted. Images were cropped to the region of interest and occasionally black triangles were added to the edges of an image to ensure the image background was a rectangular box. In Figure 2, a black box on the side of the image was used to cover a root adjacent to the root of interest. Tests of statistical significance were performed in RStudio (Version 1.1.463). Parametric tests were used when data was normal; non-parametric tests were used when data was not normal.

### Quantitative PCR Analysis

Total RNA was extracted from whole roots of five-day-old seedlings using RNeasy-Plant mini-kit (QIAGEN). Poly(dT) cDNA was prepared with Invitrogen^™^ Superscript^™^ II reverse transcriptase kit (Invitrogen), and primer efficiency was measured using a cDNA dilution series. The expression level of *ARF10* and *16* was quantified using primers in Supplemental Table 1. All individual reactions were performed in triplicates, with three biological replicates. Expression levels were normalized using ubiquitin (*UBQ*) gene expression (Supplemental Table 1). Relative expression was calculated using the delta-delta Ct method (Livak and Schmittgen, 2001).

### GUS staining

Seedlings containing *pNLP7:GUS* were grown on MS media plates for four days and transferred to MS media supplemented with 0.3 μM NAA for 24 hours. Seedlings were subsequently harvested in ice cold 95% acetone. After removing 95% acetone, GUS staining buffer (100mM NaPO4, 0.2M K3CN6, 0.2M K4CN6, 0.1%, Triton X-100) without X-gluc was vacuum infiltrated for 20 minutes at room temperature. GUS staining buffer was removed from the samples, and samples were vacuum infiltrated GUS staining buffer again with X-gluc for 20 minutes at room temperature. After the second vacuum infiltration, samples were kept overnight at 37°C. Images were captured on an Olympus (DP80) under 60X magnification using an Olympus DP80 camera.

### Accession Numbers

Arabidopsis gene IDs for the genes in this article are: At4g24020 (NLP7), At1g23080 (PIN7), At1g70940 (PIN3), AT3G11260 (WOX5), At1g76520 (PILS3), At3g20840 (PLT1), At1g51190 (PLT2), At2g28350 (ARF10), and At4g30080 (ARF16).

## ACKNOWLEDGEMENTS

This work was funded by the National Science Foundation grant number 1656392 to AIP. We thank members of the Iyer-Pascuzzi lab for comments on the manuscript.

## SUPPLEMENTAL DATA

### Supplemental Figures

**Supplemental Figure S1:** NLP7 does not impact expression of PIN3 auxin carrier in the root cap. **A and B.** *pPIN3:PIN3:GFP* in the root cap of Col −0 (A) and *nlp7-1* (B) plants on N-complete media. **C.** GFP intensity does not significantly change in the absence of *NLP7*. Black dots indicate plant samples (n > 20 for each genotype). Scale bar is 20 μm; magnification is 40X.

**Supplemental Figure S2:** In the absence of nitrate, *pPILS3:PILS3:GFP* expression decreased in wild type Col-0 plants. **A and B.** *pPILS3:PILS3:GFP* expression on N-complete (A) and NO_3_^-^ - free media (B). **C.** GFP fluorescence intensity decreased significantly in NO_3_^-^ - free media compared to N-complete media. Black dots indicate the number of plants (n = 13) used for each genotype and treatment. Three independent biological replicates were performed; representative experiment is shown. Scale bar is 20 μm; magnification is 40X. Asterisks indicates a significant difference with a two-tailed t-test (*P < 0.001).

**Supplemental Figure S3:** Auxin responses (observed through *DR5:GFP*) in the upper tiers of the columella (all tiers except the last layer) do not differ between wild type Col-0 and *nlp7-1*.Dots indicate the number of plants.

**Supplemental Figure S4:** Expression of *ARF10/16* does not change significantly between roots of Col-0 and *nlp7-1*. Expression is relative to the endogenous control gene *UBQ10*.

### Supplemental Tables

**Supplemental Table 1: Primers for qPCR and genotyping**

