## Supplemental Data for "The NIN-LIKE PROTEIN 7 (NLP7) transcription factor modulates auxin pathways to regulate root cap development"

**Supplemental Figures**

**Supplemental Figure S1**: NLP7 does not impact expression of PIN3 auxin carrier in the root cap. **A and B.** *pPIN3:PIN3:GFP* in the root cap of Col -0 (A) and *nlp7-1* (B) plants on N-complete media. **C.** GFP intensity does not significantly change in the absence of *NLP7*. Black dots indicate plant samples (n > 20 for each genotype). Scale bar is 20 µm; magnification is 40X.

**Supplemental Figure S2**: In the absence of nitrate, *pPILS3:PILS3:GFP* expression decreased in wild type Col-0 plants. **A and B.** *pPILS3:PILS3:GFP* expression on N-complete (A) and $NO_3^-$ - free media (B). **C.** GFP fluorescence intensity decreased significantly in $NO_3^-$ - free media compared to N-complete media. Black dots indicate the number of plants (n = 13) used for each genotype and treatment. Three independent biological replicates were performed; representative experiment is shown. Scale bar is 20 µm; magnification is 40X. Asterisks indicates a significant difference with a two- tailed t-test (*P < 0.001).

**Supplemental Figure S3**: Auxin responses (observed through *DR5:GFP*) in the upper tiers of the columella (all tiers except the last layer) do not differ between wild type Col-0 and *nlp7-1*. Dots indicate the number of plants.

**Supplemental Figure S4:** Expression of *ARF10/16* does not change significantly between roots of Col-0 and *nlp7-1*. Expression is relative to the endogenous control gene *UBQ10*.

**Supplemental Tables**

**Supplemental Table 1: Primers for qPCR and genotyping**

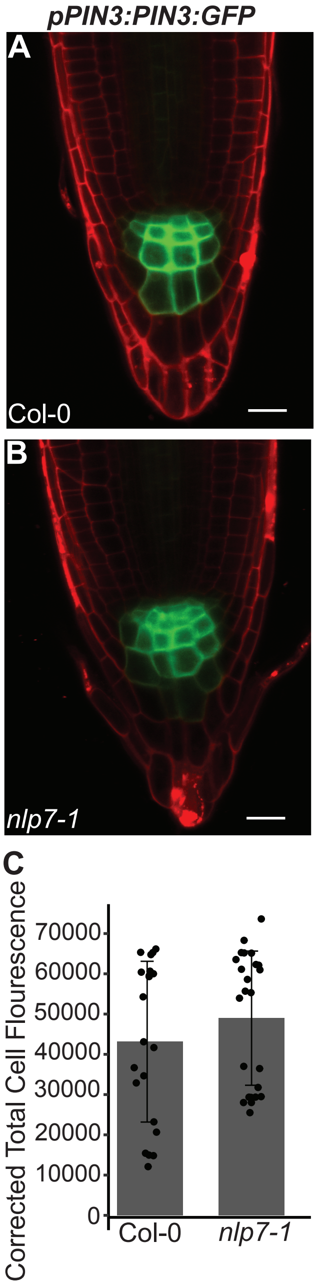


Supplemental Figure S1: NLP7 does not impact expression of PIN3 auxin carrier in the root cap. A and B. *pPIN3:PIN3:GFP* in the root cap of Col -0 (A) and *nlp7-1* (B) plants on N-complete media. C. GFP intensity does not significantly change in the absence of *NLP7*. Black dots indicate plant samples (n > 20 for each genotype). Scale bar is 20 µm; magnification is 40X.

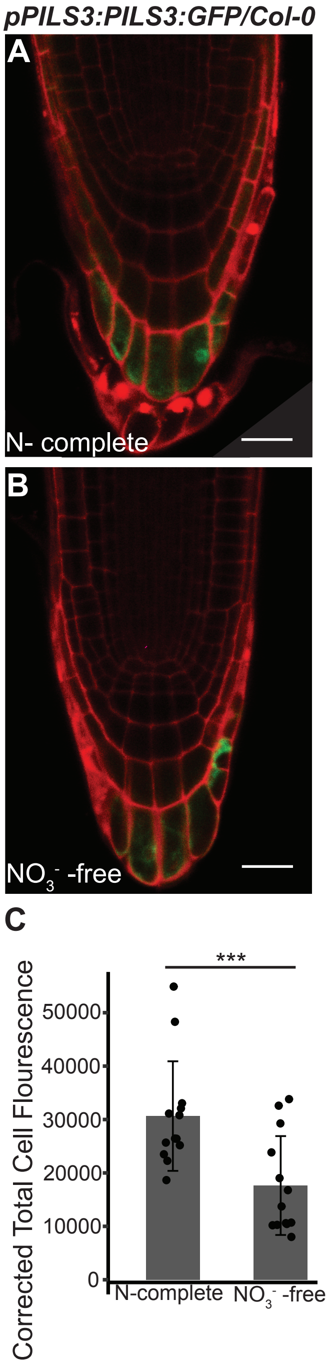


Supplemental Figure S2: In the absence of nitrate, *pPILS3:PILS3:GFP* expression decreased in wild type Col-0 plants. A and B. *pPILS3:PILS3:GFP* expression on N-complete (A) and $NO_3^-$ - free media (B). C. GFP fluorescence intensity decreased significantly in $NO_3^-$ - free media compared to N-complete media. Black dots indicate the number of plants (n = 13) used for each genotype and treatment. Three independent biological replicates were performed; representative experiment is shown. Scale bar is 20 μm; magnification is 40X. Asterisks indicates a significant difference with a two-tailed t-test (*P < 0.001).

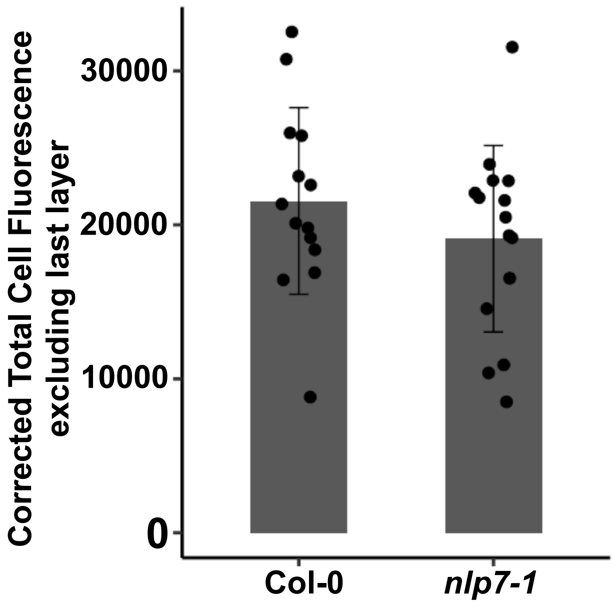


**Supplemental Figure S3**: Auxin responses (observed through *DR5:GFP*) in the upper tiers of the columella (all tiers except the last layer) do not differ between wild type Col-0 and *nlp7-1*. Dots indicate the number of plants.

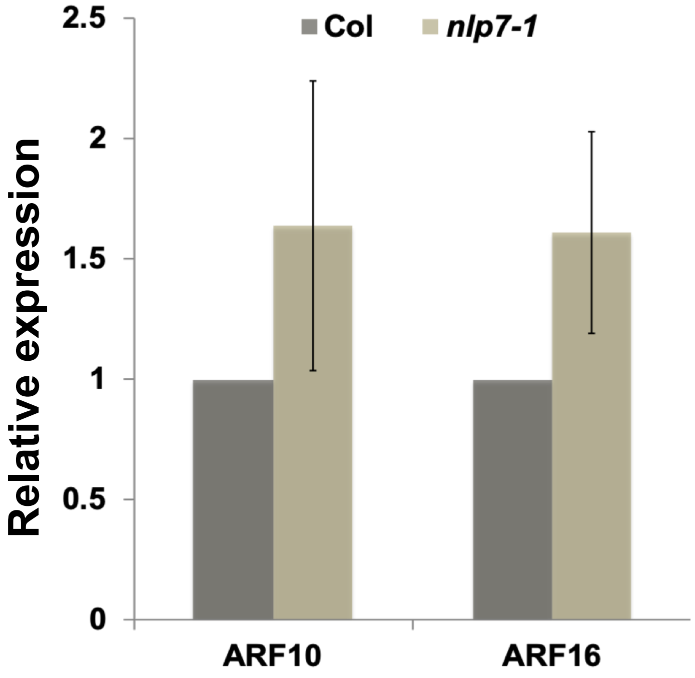


**Supplemental Figure S4:** Expression of *Auxin Responsive Factor10/16* does not change significantly between roots of Col-0 and *nlp7-1*. Expression is relative to *ubiquitin 10* (*UBQ10*).
